## Supplementary Information for "Quantitative pathogenicity and host adaptation in a fungal plant pathogen revealed by whole-genome sequencing"

### Supplementary Figures

**Supplementary Fig. 1 |** Percentage of genetic variance explained by principal component analysis.

**Supplementary Fig. 2 |** Heatmap on the kinship matrix calculated with the full SNP matrix using VanRaden method.

**Supplementary Fig. 3 |** Phenotypic distribution of PLACP and PLACN quantitative pathogenicity traits divided over cultivars from which isolates originated.

**Supplementary Fig. 4 |** Quantile-quantile plots of multi-locus mixed linear model (pink curve) and single-locus linear model (blue curve) for the different trait-cultivar associations.

**Supplementary Fig. 5 |** PLACP of individual *Zt_6_00682* mutants complemented with the virulent allele (grey boxes) and the avirulent allele (white boxes) in the *Zt_6_00682* knockout background.

**Supplementary Fig. 6 |** PLACP produced by the wild-type strain 3D7, *Zt_6_682* knockout mutants and the complementation mutants carrying the virulent (3D7) and avirulent (3D1) allele in different cultivars.

**Supplementary Fig. 7 |** PLACP of individual *Zt_3_00467* mutants on the cultivar 'Arina' (*Stb6* and *Stb15*) expressing the avirulent allele (white boxes), the virulent allele (green boxes) and the wild-type strain 3D7.

**Supplementary Fig. 8 |** PLACP of *Zt_3_00467* mutants on the cultivar 'Riband' (*Stb15*) expressing the avirulent allele (green box), the virulent allele (blue box) and the wild-type strain 3D7.

**Supplementary Fig. 9 |** Phylogenetic tree of *Zt_3_00467* and *Zt_6_00682* generated from the protein sequences of the global *Z. tritici* isolates (n=241 and 248, respectively).

### Supplementary Tables

**Supplementary Table 1 |** Descriptive statistics for quantitative pathogenicity traits assessed on differential wheat cultivars.

**Supplementary Table 2 |** Analysis of variance summary table showing the influence of isolate, cultivar and their interaction.

**Supplementary Table 3 |** Wheat differentials used to pathotype *Z. tritici* isolates and their known Stb resistance genes indicated in grey (Brown et al., 2015).

**Supplementary Table 4 |** Functional annotation of candidate pathogenicity genes.

**Supplementary Table 5 |** Diversity statistics of pathogenicity genes.

**Supplementary Table 6 |** Summary of recombination and selection analyses of pathogenicity genes.

**Supplementary Table 7 |** List of primers used for functional studies.

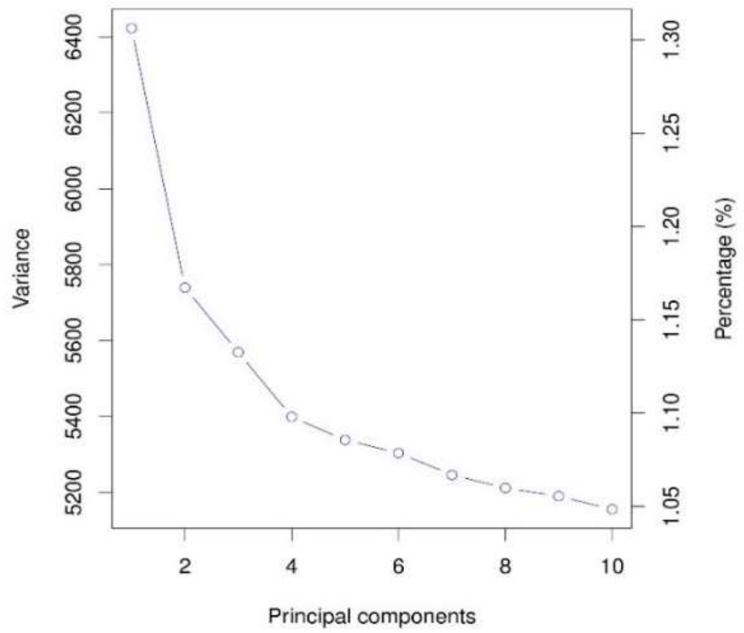


**Supplementary Fig. 1 |** Percentage of genetic variance explained by principal component analysis.

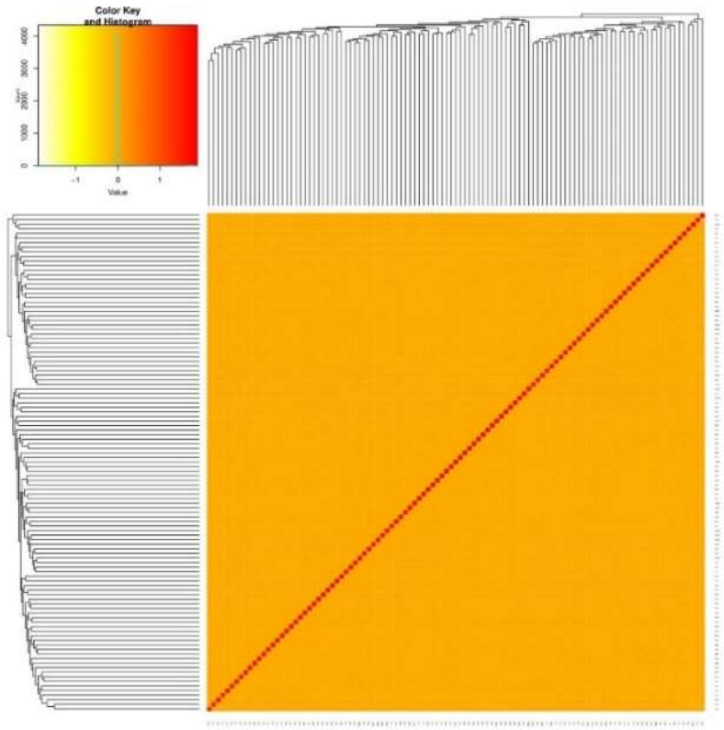


**Supplementary Fig. 2 |** Heatmap on the kinship matrix calculated with the full SNP matrix using VanRaden method.

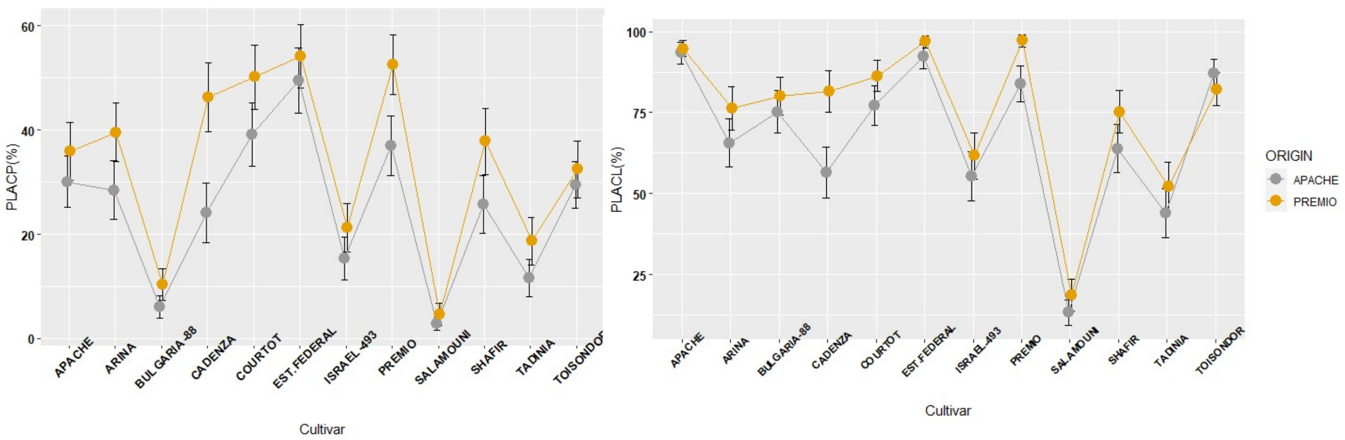


**Supplementary Fig. 3 |** Phenotypic distribution of PLACP and PLACN quantitative pathogenicity traits divided over cultivars from which isolates originated.

### PLACP

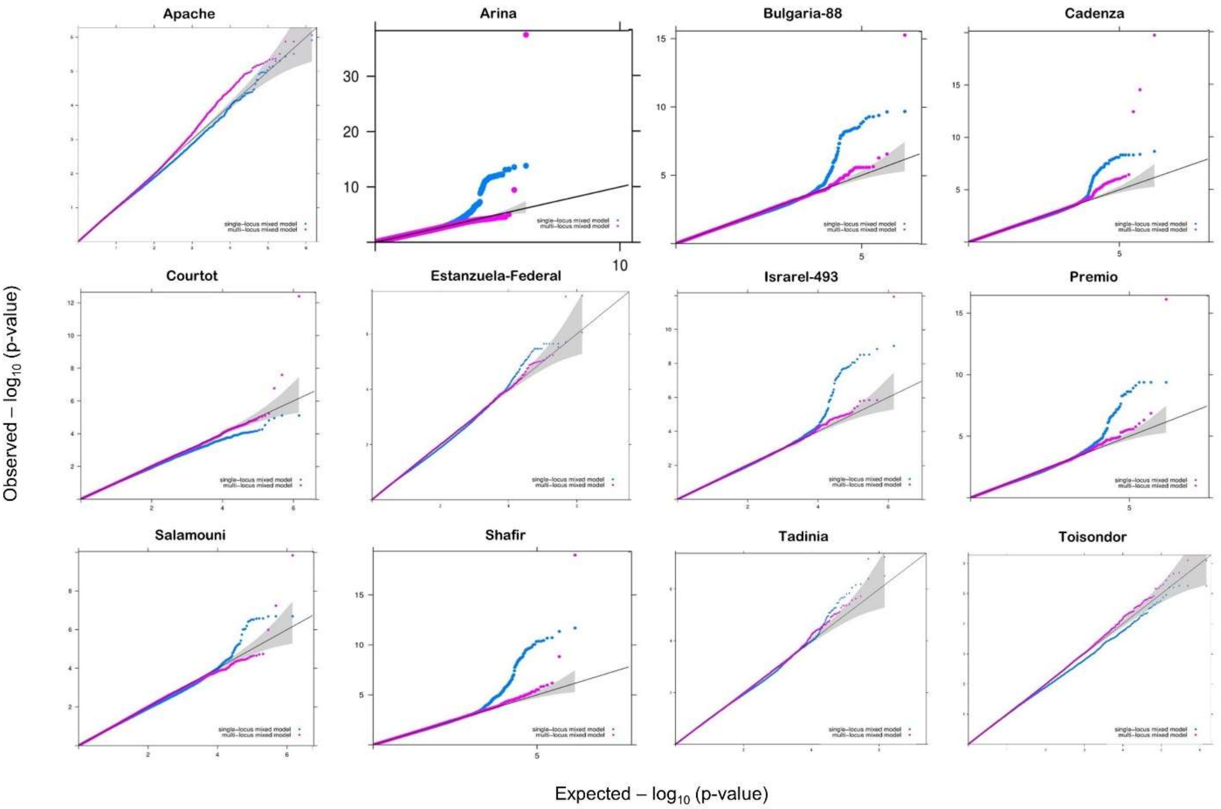


### PLACN

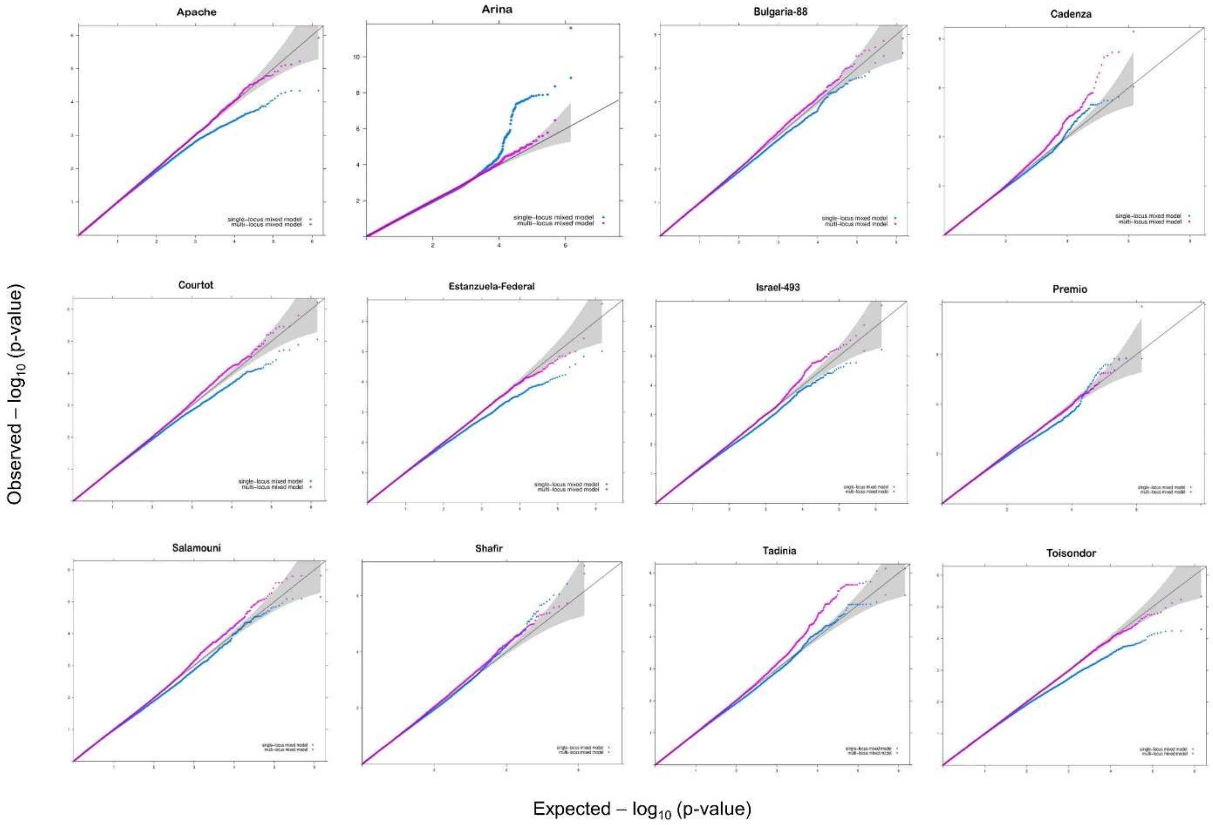


**Supplementary Fig. 4 |** Quantile-quantile plots of multi-locus mixed linear model (pink curve) and single-locus linear model (blue curve) for the different trait-cultivar associations.

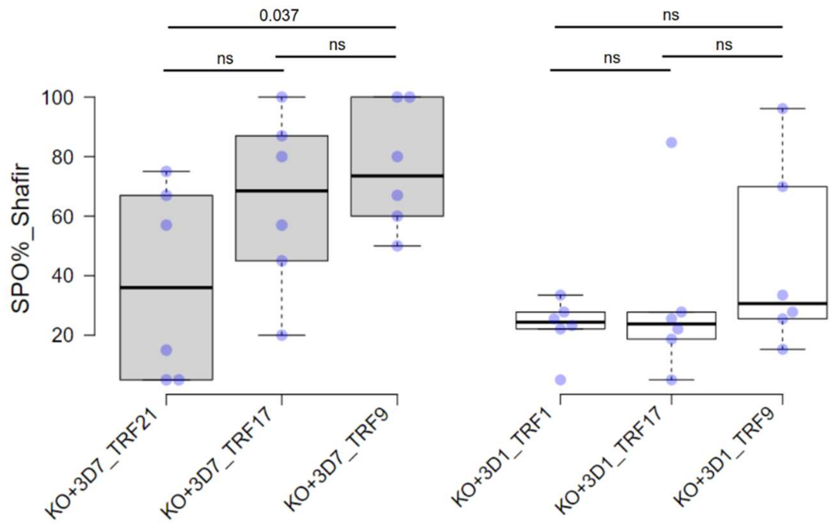


**Supplementary Fig. 5 |** PLACP of individual *Zt_6_00682* mutants complemented with the virulent allele (grey boxes) and the avirulent allele (white boxes) in the *Zt_6_00682* knockout background. Significance was determined with Wilcoxon rank-sum test. Data are presented as box plots (centre line at the median, upper bound at 75th percentile, lower bound at 25th percentile) with whiskers chosen to show the 1.5 of the interquartile range. Each dot represents one inoculated leaf.

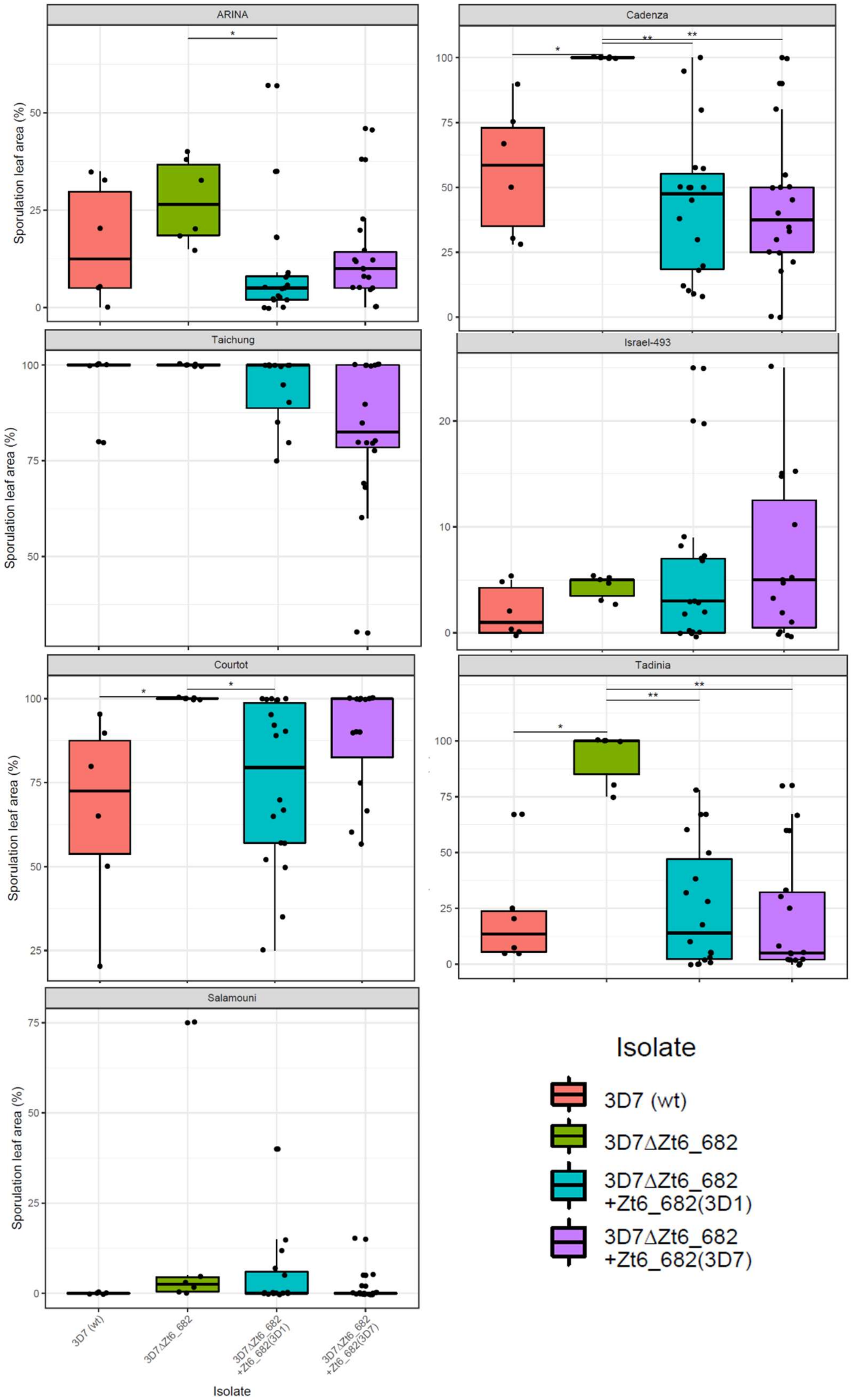


**Supplementary Fig. 6 |** PLACP produced by the wild-type strain 3D7, *Zt_6_00682* knockout mutants and the complementation mutants carrying the virulent (3D7) and avirulent (3D1) allele in different cultivars. KO mutants were highly aggressive in the cultivars ‘Cadenza’, ‘Tadinia’ and ‘Courtot’. Significance was determined with Wilcoxon rank-sum test ($P$<0.05*; $P$<0.01**). Data are presented as box plots (centre line at the median, upper bound at 75th percentile, lower bound at 25th percentile) with whiskers chosen to show the 1.5 of the interquartile range. Each dot represents one inoculated leaf.

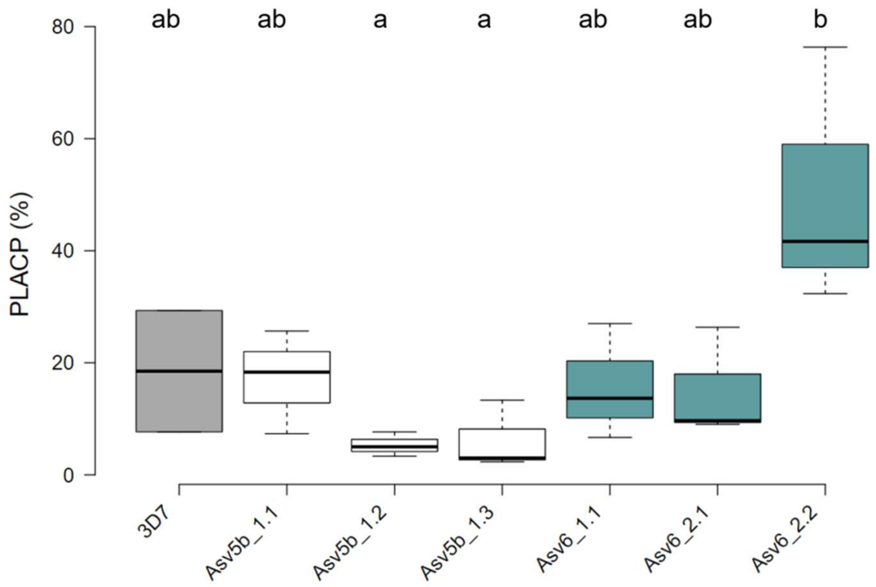


**Supplementary Fig. 7 |** PLACP of individual *Zt_3_00467* mutants on the cultivar 'Arina' (Stb6 and Stb15) expressing the avirulent allele (white boxes), the virulent allele (green boxes) and the wild-type strain 3D7. Statistical significance was determined using a Tukey's HSD test. Data are presented as box plots (centre line at the median, upper bound at 75th percentile, lower bound at 25th percentile) with whiskers chosen to show the 1.5 of the interquartile range.

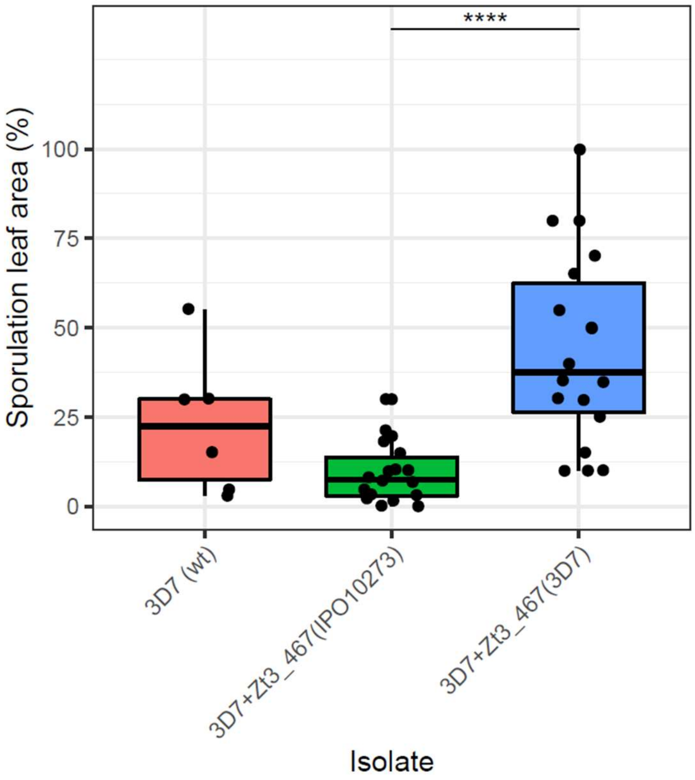


**Supplementary Fig. 8 |** PLACP of *Zt_3_00467* mutants on the cultivar 'Riband' (Stb15) expressing the avirulent allele (green box), the virulent allele (blue box) and the wild-type strain 3D7. Significance was determined with Wilcoxon rank-sum test ($P$<0.0001****). Data are presented as box plots (centre line at the median, upper bound at 75th percentile, lower bound at 25th percentile) with whiskers chosen to show the 1.5 of the interquartile range. Each dot represents one inoculated leaf.

*Zt_3_00467*

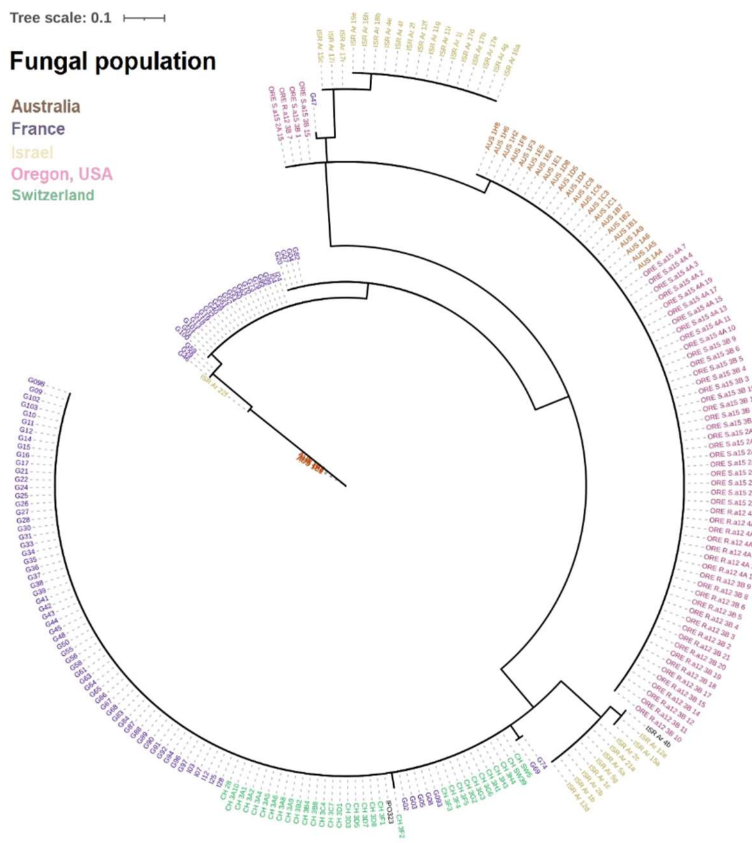


*Zt_6_00682*

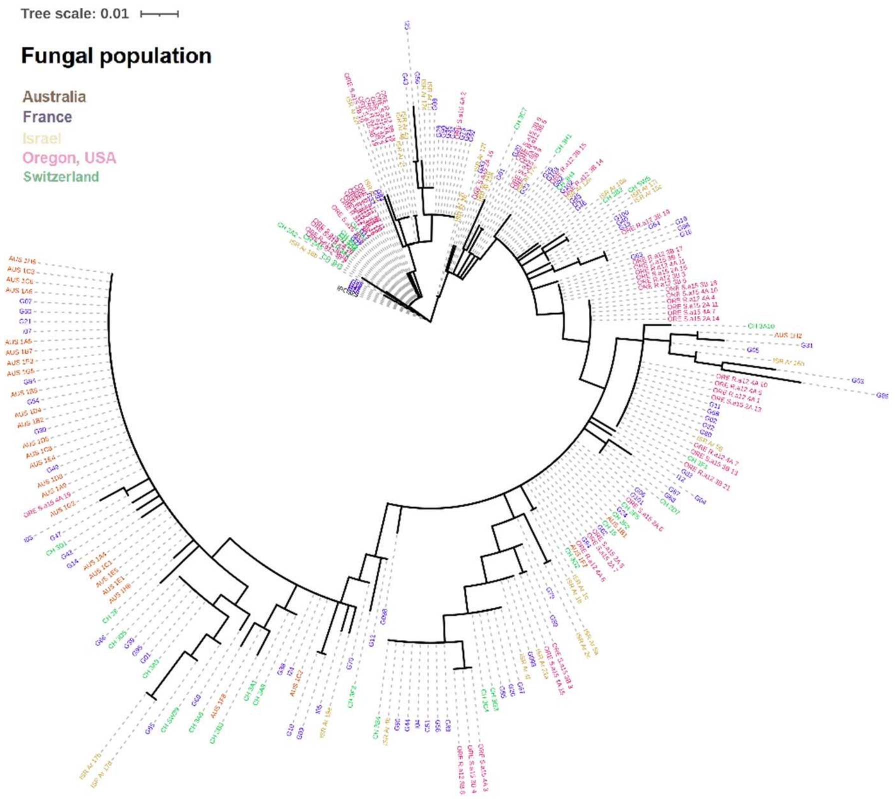


**Supplementary Fig. 9 |** Phylogenetic tree of *Zt_3_00467* (top) and *Zt_6_00682* (bottom) generated from the protein sequences of the global *Z. tritici* isolates (n=241 and 248, respectively).

**Supplementary Table 1 |** Descriptive statistics for quantitative pathogenicity traits assessed on differential wheat cultivars.

| | Range (%) | | Mean (%) | | Strandad Deviation | | Coefficient of Variation | |
|---|---|---|---|---|---|---|---|---|
| **Cultivar** | **PLACN** | **PLACP** | **PLACN** | **PLACP** | **PLACN** | **PLACP** | **PLACN** | **PLACP** |
| Apache | 35 - 100 | 0.6 - 91.8 | 93,32 | 33,74 | 11,11 | 18,28 | 11,91 | 54,18 |
| Arina | 0 - 100 | 0 - 94.7 | 73,94 | 37,1 | 36,01 | 26,9 | 48,7 | 72,51 |
| Bulgaria-88 | 0.3 - 100 | 0 - 54.1 | 76,12 | 8,9 | 28,91 | 10,85 | 37,98 | 121,94 |
| Cadenza | 0 - 100 | 0 - 94.7 | 69,87 | 37,24 | 39,61 | 31,42 | 56,69 | 84,37 |
| Courtot | 11.1 - 100 | 0 - 99.4 | 79,61 | 45,33 | 24,57 | 28,61 | 30,86 | 63,11 |
| Est-Federal | 11.1 - 100 | 1.1 - 96.4 | 93,15 | 51,68 | 16,55 | 25,88 | 17,77 | 50,07 |
| Israel-493 | 0 - 100 | 0 -74.2 | 56,5 | 18 | 30,57 | 16,89 | 54,11 | 93,81 |
| Premio | 14 - 100 | 0.1 - 93.7 | 89,64 | 44,55 | 20,9 | 24,92 | 23,31 | 55,93 |
| Salamouni | 0 - 68.3 | 0 - 35.8 | 15,43 | 3,82 | 15,78 | 5,88 | 102,29 | 154 |
| Shafir | 0 - 100 | 0 - 91.7 | 69,73 | 32,86 | 34,85 | 25,99 | 49,97 | 79,09 |
| Tadinia | 0 - 100 | 0 - 71.1 | 47,31 | 14,69 | 35,92 | 17,39 | 75,93 | 118,34 |
| Toisondor | 25 - 100 | 0 - 70 | 84,89 | 31,24 | 15,65 | 18,08 | 18,44 | 57,88 |

**Supplementary Table 2 |** Analysis of variance summary table showing the influence of isolate, cultivar and their interaction.

| | PLACP | | | | | PLACN | | | |
|---|---|---|---|---|---|---|---|---|---|
| | ***df*** | **MS** | ***F*** | ***P*** | | ***df*** | **MS** | ***F*** | ***P*** |
| **Isolate** | 108 | 6266 | 10,73 | *** | | 108 | 11027 | 18,83 | *** |
| **Cultivar** | 11 | 57501 | 98,52 | *** | | 11 | 128974 | 220,27 | *** |
| **Isolate x Cultivar** | 1188 | 1016 | 1,74 | *** | | 1188 | 1456 | 2,48 | *** |
| **Error** | 2045 | 584 | | | | 2045 | 586 | | |

**Supplementary Table 3 |** Wheat differentials used to pathotype *Z. tritici* isolates and their known Stb resistance genes indicated in grey (Brown *et al.*, 2015)[1].

| Cultivar/*R* gene | ***Stb1*** | ***Stb2*** | ***Stb3*** | ***Stb4*** | ***Stb5*** | ***Stb6*** | ***Stb7*** | ***Stb8*** | ***Stb9*** | ***Stb10*** | ***Stb11*** | ***Stb12*** | ***Stb13*** | ***Stb14*** | ***Stb15*** | ***Stb16q*** | ***StbSm3*** | **Reference** |
|---|---|---|---|---|---|---|---|---|---|---|---|---|---|---|---|---|---|---|
| Bulgaria-88 | ■ | | | | | ■ | | | | | | | | | | | | Adhikari *et al.* 2004c[2]; Chartrain *et al.* 2005b[3] |
| Veranopolis | | ■ | | | | ■ | | | | | | | | | | | | Adhikari *et al.* 2004b[4]; Chartrain *et al.* 2005b[3] |
| Israel-493 | | | ■ | | | ■ | | | | | | | | | | | | Adhikari *et al.* 2004b[4]; Chartrain *et al.* 2005b[3] |
| Tadinia | | | | ■ | | ■ | | | | | | | | | | | | Adhikari *et al.* 2004a[5]; Chartrain *et al.* 2005b[3]; Somasco *et al.* 1996[6] |
| CS-synthetic (69)7D | | | | | ■ | ■ | | | | | | | | | | | | Arraiano *et al.* 2001[7] |
| Cadenza | | | | | | ■ | | | ?[1] | | | | | | | | | Zhong *et al.* 2017[8]; Saintenac *et al.* 2018[9] |
| Shafir | | | | | | ■ | | | | | | | | | | | | Brading *et al.* 2002[10] |
| Est.-Federal | | | | | | | ■ | | | | | | | | | | | McCartney *et al.* 2003[11] |
| M6-synthetic (W-7984) | | | | | | | | ■ | | | | | | | | | | Adhikari *et al.* 2003[12] |
| Courtot | | | | | | | | | ■ | | | | | | | | | Chartrain *et al.* 2009[13]; Amezrou *et al.* 2023[14] |
| Kavkaz-K4500 | | | | | | ■ | ■ | | | ■ | | ■ | | | | | | Chartrain *et al.* 2005a[15] |
| TE-9111 | | | | | | ■ | ■ | | | | ■ | | | | | | | Chartrain *et al.* 2005c[16] |
| Apache | | | | ? | ? | | | | | | ■ | | | | | | | Ghaffary *et al.* 2011[17] |
| Salamouni | | | | | | | | | | | | | ■ | ■ | | | ■ | Cowling 2006[18] |
| Arina | | | | | | ■ | | | | | | | | | ■ | | | Arraiano *et al.* 2007[19]; Chartrain *et al.* 2005b[3] |
| M3-synthetic (W-7976) | | | | | | | | | | | | | | | | ■ | | Ghaffary *et al.* 2012[20]; Saintenac et al. 2021[21] |

[1] *Stb9* was detected in the cultivar Cadenza for the first time in this study

**Supplementary Table 4 |** Functional annotation of candidate pathogenicity genes.

| Gene name | Gene ID | Protein length | % of Cysteines | SignalP (probability) | # of predicted TMHs | EffectorP (probability) | Conserved protein domain | | |
|---|---|---|---|---|---|---|---|---|---|
| | | | | | | | Domain | E-value | Description |
| | *Zt_1_00196* | 422 | 2,61 | | 5 | | | | |
| *AvrStb9* | *Zt_1_00693* | 727 | 1,13 | 0,99 | | | Peptidase_S41 superfamily | 3,99E-02 | Peptidase_S41 |
| | *Zt_1_01325* | 95 | 0,00 | | 2 | | DPM2 | 6,29E-32 | dolichol phosphate-mannose biosynthesis regulatory |
| | *Zt_1_01674* | 408 | 1,23 | 1,00 | | | TrxB | 6,10E-21 | Thioredoxin reductase |
| | *Zt_1_01987* | 475 | 0,63 | | | | cupin_RmlC-like | 0,00 | cupin_like |
| *MGAMN10* | *Zt_2_00163* | 506 | 0,79 | 1,00 | | | Glyco_hydro_47 | 0,00 | Glycosyl hydrolase family 47 |
| | *Zt_2_00165* | 430 | 0,00 | 1,00 | | | Ish1 | 1,53E-10 | Putative stress-responsive nuclear envelope protein |
| | *Zt_2_01230* | 96 | 0,00 | 1,00 | | 0,73 | | | |
| | *Zt_3_00467* | 126 | 6,35 | 1,00 | | 0,86 | | | *SSP - Z. tritici specific* |
| *AvrStb6* | - | 83 | 14,63 | 1,00 | | 0,96 | | | *SSP - Z. tritici specific* |
| | *Zt_5_00252* | 385 | 1,04 | | 6 | | | | |
| | *Zt_6_00224* | 69 | 8,70 | 1,00 | | 0,85 | | | *SSP - Z. tritici specific* |
| | *Zt_6_00425* | 329 | 0,91 | | | | | | |
| | *Zt_6_00682* | 305 | 0,33 | | | 0,50 | Methyltransf_25 | 5,33E-16 | SAM-dependent methyltransferase |
| | *Mycgr3G110052* | 179 | 1,12 | 0,99 | | 0,58 | | | *SSP - Z. tritici specific* |
| | *Zt_9_00069* | 249 | 0,00 | 0,87 | | | adh_short | 1,56E-31 | Short chain dehydrogenase |
| | *Zt_13_00071* | 444 | 1,13 | | | | | | |
| | *Mycgr3G95574* | 114 | 4,39 | 1,00 | | 0,60 | | | *SSP - Z. tritici specific* |
| | *Mycgr3G106276* | 109 | 4,59 | | | | | | *Z. tritici specific* |

Supplementary Table 4. Continued.

| Gene name | Gene ID | PHIB-blast | | | | | |
|---|---|---|---|---|---|---|---|
| | | PHI-base ID | Gene | E-value | % Identi-ties | pathogen species | Phenotype |
| | *Zt_1_00196* | 5822 | Fgsg1989 | 8,94E-34 | 31,35 | *Fusarium graminearum* | unaffected pathogenicity |
| *AvrStb9* | *Zt_1_00693* | | | | | | |
| | *Zt_1_01325* | | | | | | |
| | *Zt_1_01674* | 6471 | Trr2 | 2,41E-17 | 25,95 | *Beauveria bassiana* | reduced virulence |
| | *Zt_1_01987* | 542 | hmgA | 1,39E-142 | 52,86 | *Pseudomonas aeruginosa* | increased virulence |
| *MGAMN10* | *Zt_2_00163* | 251 | msdS/Af msdC | 0,00 | 59,10 | *Aspergillus fumigatus* | unaffected pathogenicity |
| | *Zt_2_00165* | | | | | | |
| | *Zt_2_01230* | | | | | | |
| | *Zt_3_00467* | | | | | | |
| *AvrStb6* | - | | | | | | |
| | *Zt_5_00252* | 587 | Fgsg4159 | 4,43E-28 | 31,27 | *Fusarium graminearum* | unaffected pathogenicity |
| | *Zt_6_00224* | | | | | | |
| | *Zt_6_00425* | | | | | | |
| | *Zt_6_00682* | 3881 | LaeA | 4,74E-10 | 24,62 | *Aspergillus flavus* | reduced virulence |
| | *Mycgr3G110052* | | | | | | |
| | *Zt_9_00069* | 1992 | GzZC37 | 3,42E-48 | 55,53 | *Fusarium graminearum* | unaffected pathogenicity |
| | *Zt_13_00071* | | | | | | |
| | *Mycgr3G95574* | | | | | | |
| | *Mycgr3G106276* | | | | | | |

**Supplementary Table 5 |** Diversity statistics of pathogenicity genes.

| | Gene presence (%) | Segr. Sites | Haplo-types | Haplotype diversity | π | Tajima D | | Fu & Li D | | Fu & Li F | | ZnS (LD) | GC content (%) | TE distance (Kb) |
|---|---|---|---|---|---|---|---|---|---|---|---|---|---|---|
| *Mean* | 97,04 | 104,4561 | 47,737 | 0,8376 | 0,0196 | -0,0340 | | -1,1770 | | -0,7986 | | 0,1546 | 0,5425 | 15,25 |
| *Median* | 100,00 | 65,00 | 39,00 | 0,90 | 0,02 | 0,88 | | -0,26 | | 0,04 | | 0,16 | 0,55 | 4,15 |
| *Avrstb6* | 100,00 | 65 | 18 | 0,7750 | 0,0255 | -1,2017 | n.s. | 0,5553 | n.s. | -0,2092 | n.s. | 0,3104 | 0,5448 | 2,07 |
| *AvrStb9* | 100,00 | 81 | 28 | 0,8362 | 0,0051 | -0,9460 | n.s. | 0,4286 | n.s. | -0,1746 | n.s. | 0,1629 | 0,5392 | 2,93 |
| *Mycgr3G106276* | 100,00 | 28 | 35 | 0,9086 | 0,0230 | 1,0792 | n.s. | 0,5038 | n.s. | 0,8379 | n.s. | 0,1615 | 0,5146 | 4,15 |
| *Mycgr3G110052* | 51,46 | 145 | 10 | 0,5806 | 0,1182 | 1,2962 | n.s. | 2,0417 | ** | 2,0673 | ** | 0,5112 | 0,4457 | 2,23 |
| *Mycgr3G95574* | 100,00 | 48 | 40 | 0,9098 | 0,0323 | -0,6859 | n.s. | 1,3557 | n.s. | 0,6459 | n.s. | 0,1696 | 0,5904 | 3,55 |
| *Zt_1_00196* | 100,00 | 31 | 29 | 0,6674 | 0,0015 | -2,0907 | * | -5,5494 | ** | -4,8307 | ** | 0,0355 | 0,5492 | 0,48 |
| *Zt_1_01325* | 100,00 | 17 | 13 | 0,5189 | 0,0108 | 0,8775 | n.s. | -1,3548 | n.s. | -0,6079 | n.s. | 0,3324 | 0,5492 | 2,98 |
| *Zt_1_01674* | 100,00 | 75 | 68 | 0,9743 | 0,0105 | -0,1741 | n.s. | -1,9473 | # | -1,3726 | n.s. | 0,0510 | 0,5487 | 14,59 |
| *Zt_1_01987* | 100,00 | 125 | 55 | 0,9271 | 0,0300 | 2,2148 | * | 1,1640 | n.s. | 1,9050 | * | 0,1187 | 0,5816 | 17,88 |
| *Zt_13_00071* | 98,06 | 96 | 43 | 0,9061 | 0,0293 | 1,4282 | n.s. | -1,0453 | n.s. | 0,0431 | n.s. | 0,1627 | 0,5508 | 16,55 |
| *Zt_2_00163* | 100,00 | 145 | 92 | 1,0000 | 0,0322 | 2,4158 | * | 0,7520 | n.s. | 1,7482 | * | 0,0534 | 0,5541 | 21,96 |
| *Zt_2_00165* | 100,00 | 50 | 45 | 0,8247 | 0,0133 | -0,1118 | n.s. | -2,1126 | n.s. | -1,5075 | n.s. | 0,0830 | 0,5761 | 17,21 |
| *Zt_2_01230* | 100,00 | 17 | 22 | 0,8093 | 0,0047 | -1,1630 | n.s. | -4,3619 | ** | -3,6519 | ** | 0,0199 | 0,5396 | 0,18 |
| *Zt_3_00467* | 97,09 | 33 | 16 | 0,8358 | 0,0145 | 2,4201 | * | 1,0260 | n.s. | 1,9408 | # | 0,3217 | 0,5489 | 0,04 |
| *Zt_5_00252* | 100,00 | 295 | 52 | 0,9044 | 0,0520 | 1,7580 | # | 1,2515 | n.s. | 1,7403 | * | 0,2805 | 0,5343 | 36,53 |
| *Zt_6_00224* | 99,03 | 22 | 24 | 0,8515 | 0,0050 | -0,5148 | n.s. | -2,7348 | ** | -2,1360 | * | 0,0513 | 0,5568 | 5,19 |
| *Zt_6_00425* | 100,00 | 156 | 97 | 0,9989 | 0,0495 | 1,0488 | n.s. | -0,2640 | n.s. | 0,4079 | n.s. | 0,0906 | 0,5856 | 10,19 |
| *Zt_6_00682* | 100,00 | 43 | 39 | 0,9299 | 0,0212 | 1,1329 | n.s. | -0,6695 | n.s. | 0,0652 | n.s. | 0,2225 | 0,6068 | 4,39 |
| *Zt_9_00069* | 98,06 | 95 | 51 | 0,8980 | 0,0128 | -1,6910 | # | -2,5183 | * | -2,4677 | * | 0,1406 | 0,5606 | 7,13 |

*n.s. not significant*

*# P<0.1*

** P<0.05*

*** P<0.01*

**** P<0.001*

**Supplementary Table 6 |** Summary of recombination and selection analyses of pathogenicity genes.

| | Recombination | | codeML | | | | | | | |
|---|---|---|---|---|---|---|---|---|---|---|
| | | | | | M8 model parameters | | | | | |
| Gene ID | Breakpoints[a] | Events[b] | M1a vs M2a | M7 vs M8 | dN/dS[c] | lnL[d] | ω[e] | p[f] | PSS[g] | % of positively selected sites |
| *AvrStb6* | 2 | - | <.001 | <.001 | 2,84 | -726,623 | 17,129 | 0,147 | 10 (9) | 10,98 |
| *AvrStb9* | 2 | 1 | <.001 | <.001 | 1,04 | -3137,804 | 15,137 | 0,069 | 12 (5) | 0,69 |
| *Mycgr3G106276* | 1 | - | <.001 | <.001 | 1,18 | -775,735 | 10,420 | 0,092 | 6 (5) | 4,59 |
| *Mycgr3G110052* | 2 | 2 | Presence/absence polymorphism + partial deletion | | | | | | | NA |
| *Mycgr3G95574* | - | 11 | <.001 | <.001 | 0,34 | -1243,004 | 4,879 | 0,048 | 3 (2) | 1,75 |
| *Zt_1_00196* | - | - | *NS* | *NS* | 0,12 | -1942,141 | 7,398 | 0,010 | none | NA |
| *Zt_1_01325* | - | - | *NS* | *NS* | 1,00 | -398,754 | 1,000 | 1,000 | none | NA |
| *Zt_1_01674* | 2 | - | <.001 | <.001 | 0,12 | -2623,182 | 4,828 | 0,016 | 4 (3) | 0,74 |
| *Zt_1_01987* | 5 | 17 | <.001 | <.001 | 0,04 | -5452,238 | 1,940 | 0,017 | 2 (2) | 0,42 |
| *Zt_13_00071* | 1 | 12 | <.001 | <.001 | 0,33 | -4221,104 | 2,907 | 0,084 | 9 (3) | 0,68 |
| *Zt_2_00163* | 2 | 10 | *NS* | *NS* | 0,25 | -650,262 | 1,000 | 0,000 | none | NA |
| *Zt_2_00165* | 1 | 4 | <.001 | <.001 | 0,17 | -3637,802 | 4,708 | 0,031 | 9 (7) | 1,63 |
| *Zt_2_01230* | - | - | *NS* | *NS* | 0,17 | -450,211 | 1,000 | 0,000 | none | NA |
| *Zt_3_00467* | 2 | 1 | <.001 | <.001 | 7,03 | -588,796 | 50,813 | 0,127 | 9 (6) | 4,76 |
| *Zt_5_00252* | 2 | 6 | <.001 | <.001 | 0,41 | -3131,581 | 3,366 | 0,102 | 13 (5) | 1,30 |
| *Zt_6_00224* | - | - | <.001 | <.001 | 0,87 | -488,880 | 7,113 | 0,123 | 3 (2) | 2,90 |
| *Zt_6_00425* | 2 | 12 | <.001 | <.001 | 0,31 | -8130,354 | 2,689 | 0,091 | 27 (24) | 7,29 |
| *Zt_6_00682* | 1 | 1 | <.001 | <.001 | 0,13 | -2904,540 | 7,783 | 0,010 | 3 (3) | 0,98 |
| *Zt_9_00069* | - | 2 | *NS* | *NS* | 0,11 | -1950,595 | 1,000 | 0,070 | none | NA |

[a]Number of recomination breakpoints detected by GARD

[b]Number of recombination event detected by at least 5 algorithms in RDP

[c]non-synonymous/synonymous substitution ratios across all gene tree under the model M8

[d]lnL the likelihood of the experimental model

[e]dN/dS ratio estimated in the model parameters

[f]the portion of positively selected sites estimated in the model parameters

[g]Positiveley selected sites with a NEB posterior probability > 0.95. Sites with a BEB probability > 0.95 are under parenthesis.

**Supplementary Table 7 |** List of primers used for functional studies.

| No. | Primer name | Primer sequence | Amplification |
|---|---|---|---|
| ***Zt_6_00682*** | | | |
| 1 | pNOV_for | ATGACGCGGGACAAG | Vector, KO and ectopic transformants validation |
| 2 | pNOV_rev | TAACACATTGCGGATACG | Vector, KO and ectopic transformants validation |
| 3 | Hygro_for | tgatattgaaggagcat | cloning KO transformants |
| 4 | Hygro_rev | TCTATTCCTTTGCCC | cloning KO transformants |
| 19 | Hygro_rev2 | GGGATCAGCAATCGC | KO vector validation |
| 20 | Hygro_for2 | CTGCCTGAAACCGAAC | KO vector validation |
| 23 | pNOV_inv_for | ATTCGAATCCGGAGC | cloning KO |
| 24 | pNOV_inv_rev | ATGGTCGACTCTAGAGGATC | cloning KO |
| 47 | 5’ 3D7 IPO for | GGATCCTCTAGAGTCGACCATCATGATGACACCTCAGGCGTC | cloning KO |
| 48 | 5’ 3D7 IPO rev | CCAAAAAATGCTCCTTCAATATCAGTTGAAGTCTTTTTGATGGTGGTA | cloning KO |
| 49 | 3’ 3D7 for | CGTCCGAGGGCAAAGGAATAGATGGTTGGAAGTGGTGCTGATG | cloning KO |
| 51 | 3’ 3D7 IPO rev | TGCGGCCGCTCCGGATTCGAATATCCGCGAGGAGATGGAGAATA | cloning KO |
| 72 | Gibsecto_3D7_for | GAATTCGAGCTCGGTACGTTGGTGGATGTGATCTGTG | ectopic cloning |
| 73 | Gibsecto_3D7_rev | AATGCTCCTTCAATATCAATCAAAACAGAGCTCTCGTG | ectopic cloning |
| 74 | Gibsecto_3D1_for | GAATTCGAGCTCGGTACGTTGGTGGATGTGATCTGTG | ectopic cloning |
| 75 | Gibsecto_3D1_rev | AATGCTCCTTCAATATCAACTACGAAAGTCAAAACGGA | ectopic cloning |
| 76 | Seq 3D1 3D7 for | CTAGTCCATCTCGCCATTC | validation ectopic vector and ectopic transformants |
| 77 | Seq 3D1 3D7 rev | CCCTCACCACGAAACTC | validation ectopic vector and ectopic transformants |
| 78 | Seq 3D1 3D7 for 2 | ATCCGCAAAGTCGTCG | validation vecteur ectopique |
| 79 | Seq 3D1 3D7 for 3 | CTTCAGGATGCGAGAC | validation vecteur ectopique |
| 94 | ValidKO_3D7IPO_F | TTGAGTTTCGTGGTG | Validation KO transformants |
| 95 | ValidKO_3D7IPO_R | GGTCATGCATACATTG | Validation KO transformants |
| 46 | pNOV_sulf_rev | CGCCTGGACGACTAAAC | validation ectopic vector |
| 117 | Pnov_for BIS | CAGCGGCCATTTAAATC | validation ectopic transformants |
| ***Zt_3_00467*** | | | |
| ASV10p | Zt_3_00467_AVR_IF_F1 | GCGCGCCGAATTCGAGCTCATTAACCGCTCTTGGTCGCT | Ectopic expression of Zt_3_00467 3D7 and IPO10273 alleles |
| ASV11p | Zt_3_00467_AVR_IF_R1 | CCAACATGGTGGAGTGAGGGTCCAGGAACAACCTTGCAC | Ectopic expression of Zt_3_00467 3D7 and IPO10273 alleles |
